# Do carbonated beverages reduce bleeding from gill injuries in angled Northern Pike?: In Prep for North American Journal of Fisheries Management

**DOI:** 10.1101/2020.06.15.150797

**Authors:** Alexandria Trahan, Auston Chhor, Michael J. Lawrence, Jacob W. Brownscombe, Daniel Glassman, Connor H. Reid, Alice E.I. Abrams, Andy J. Danylchuk, Steven J. Cooke

## Abstract

The premise of catch-and-release angling is that most fish survive fisheries interactions. Therefore, it is common for anglers, management agencies, and other organizations to share information on handling practices and other strategies that are believed to improve fish welfare and survival. Recent media coverage has sensationalized the use of carbonated beverages to treat bleeding fish, an intervention that is purported to stop bleeding but has yet to be validated scientifically. We captured Northern Pike (*Esox lucius*) via hook and line, experimentally injured their gills in a standardized manner, and treated them with either Mountain Dew, Coca Cola, or carbonated lake water and observed the duration and intensity of bleeding, as well as overall blood loss (using gill colour as a proxy) while the fish was held in a lake water bath. As a control, we had a group of experimentally injured fish that did not have liquid poured over their gills before the observation period. All treatments and the control were conducted at two different water temperatures (11-18 °C and 24-27 °C) to determine if the effects of pouring carbonated beverages over injured gills is temperature dependent. When compared to the control, we found that the duration and intensity of bleeding increased regardless of the type of carbonated beverages used in this study, and there was no effect of water temperature. Use of chilled versus ambient temperature beverages similarly had no influence on outcomes. As such, there is no scientific evidence to support the use of carbonated beverages for reducing or stopping blood loss for fish that have had their gills injured during recreational angling based on the context studied here. This study reinforced the need to scientifically test angler anecdotes and theories when it comes to best practices for catch-and-release fishing.

## Introduction

Recreational angling is a common practice around the globe. Although some fish are harvested, a greater percentage of them are released (Cooke and Cowx 2004). Catch-and-release (C&R) occurs when recreational anglers comply with local harvest regulations or when it is adopted voluntarily based on their conservation ethic (Arlinghaus et al. 2007). Regardless of the reason, the general premise with C&R is that most released will fish survive angling-released stress and physical injuries (Wydoski 1977). Hooking injury is the most important factor influencing whether a fish survives a C&R event, with hook injury to critical areas, such as the gills or deeply in the esophagus, yielding comparatively higher mortality than when fish are hooked in areas such as the corner of the jaw (Muoneke and Childress 1994; Bartholomew and Bohnsack 2005; Arlinghaus et al., 2007; Cooke and Schramm 2007). Based on this, there has been considerable effort to develop techniques to reduce hooking injuries in critical areas, including bleeding that can occur where the hook penetrates the fish (reviewed in Brownscombe et al. 2017).

Using carbonated beverages, such as Mountain Dew and Coca Cola, poured over gills of a fish to reduce or stop bleeding caused by hooking injury is gaining in popularity within the recreational angling community as a best practice for C&R. There is even a Facebook page, “Save a Million Fish”, where recreational anglers share videos and stories of how they have saved Muskellunge (*Esox masquinongy*) with deep hooking injuries (Anderson 2018), including pouring carbonated beverages over the gills. Popular media articles in support of this practice suggest that carbon dioxide in the beverages causes vasoconstriction of the blood vessels to slow or stop bleeding (Pyzer 2015, 2019), or that phosphoric acid in carbonated beverages causes coagulation (Green 2015; Bardin 2019). However, there is no empirical research on the effectiveness of carbonated beverages for impeding bleeding in injured fish.

Gills are a multifunctional organ for fish, playing critical roles in gas exchange, ion and water balance, ammonia excretion, and acid-base balance (Evans et al. 2005). Carbonated beverages have low pH coupled with high levels of carbon dioxide (CO_2_) in aqueous solution, various sugars, caffeine (if not caffeine free), phosphoric acid (H_3_PO_4_; Coca Cola), and citric acid (C_6_H_8_O_7_; Mountain Dew). Several of these compounds could have an effect on gill injuries. There is a rich literature describing the effects of low pH water on ion regulation, ammonia excretion, and metabolic acid (H^+^) excretion (Wood and McDonough 1988; Evans et al. 2005; Kwong et al. 2014). The elevation of external CO_2_ levels when carbonated beverages are poured over the gills may drive CO_2_ into the fish’s body by reversing the normal gradient for CO_2_ excretion from blood to water (Gilmour, 2010, 2001; Gilmour and Perry 2009). In addition, chemoreceptors that detect changes in CO_2_ are located in the gill, and may activate cardiorespiratory reflexes such as increased breathing, bradycardia, and peripheral vasoconstriction (Reid et al. 2000; Brauner et al. 2019; Tresguerres et al. 2019). Bradycardia, or slowing of heart rate, may transiently reduce bleeding (Perry and Desforges 2006). Also, caffeine is an non-specific adenosine receptor antagonist that may alter chemoreceptor signalling (Coe et al. 2017).

Regardless of the mechanism, the technique of pouring carbonated beverages over bleeding hook injuries of fish continues to be promoted, but debates as to the efficacy of its uses have also ensued (Neuharth 2019). Air exposure also occurs when carbonated beverages are poured over injured gills, which could have negative physiological effects (Cooke and Sneddon 2007). Given the growing popularity of this technique yet uncertainty related to its effectiveness, we set out to provide the first scientific evidence whether carbonated beverages should be considered a best practice for C&R because they reduce or stop bleeding from the gills of teleost fish. For this study, we angled Northern Pike (*Esox lucius*) and simulated a hooking injury to a standardized section of gill filaments. We then poured carbonated beverages over the gills (i.e., Coca Cola, Mountain Dew, or carbonated lake water), and quantified the duration and intensity of bleeding, as well as overall blood loss, in comparison to a control. Since metabolism and related blood flow in fish are positively correlated with water temperature, we also tested the effects of carbonated beverages on bleeding in Northern Pike at two different temperature regimes. Further, at the warmer water temperature we tested whether use of chilled beverages influenced outcomes relative to beverages at ambient temperatures.

## Methods

### Animal Welfare

All experiments were conducted in accordance to regulations and guidelines set by the Canadian Council on Animal Care (Carleton University protocol AUP #110558), and veterinary professionals were consulted during project development. Northern Pike Fish were collected under Scientific Collection Permit #08577 from the Ontario Ministry of Natural Resources and Forestry.

### Fish Capture

Northern Pike were selected for study owing to their relevance in addressing the question of interest and their popularity as a target recreational angling species (Paukert et al. 2001). The study was conducted on wild Norther Pike from Lake Opinicon, Ontario, Canada (44.5590° N, 76.3280° W), and all fish were capture from a boat using conventional medium-heavy rod and reel and a variety of crankbaits, chatter baits, and spinner baits. Once hooked, fish were retrieved, brought into the boat using a rubberized landing net, and immediately transferred to a water-filled trough for hook removal (underwater). Only fish that were hooked in the jaw were used in experiments to avoid confounding effects between lure-induced gill damage and experimental gill injury. In addition, fish that were bleeding from hooking site or the gills upon capture (<10% of fish) were not used in the study and were immediately released. Following hook removal, the total length (TL, mm) of the fish was recorded.

### Experimental injury and post-injury monitoring

Gill injuries were simulated by using end-cutting pliers to remove a 0.9 cm by 0.9 cm section of gill filaments from the right middle gill arch (Fig. 1). The procedure was conducted while fish were held in the water-filled trough to eliminate the air exposure. Once gills were clipped, one of three carbonated beverage treatments or the lake water control was applied (details below), and fish transferred to a white bottomed cooler (52 cm x 26.5 cm) containing nonaerated lake water (∼25 liters). When pouring carbonated beverages over the gills, a standardized volume of 150 mL was used. We also included a reference or baseline group of fish that did not have any gill filaments clipped nor liquid poured over the gills before being transferred to the cooler.

**Figure 1.**
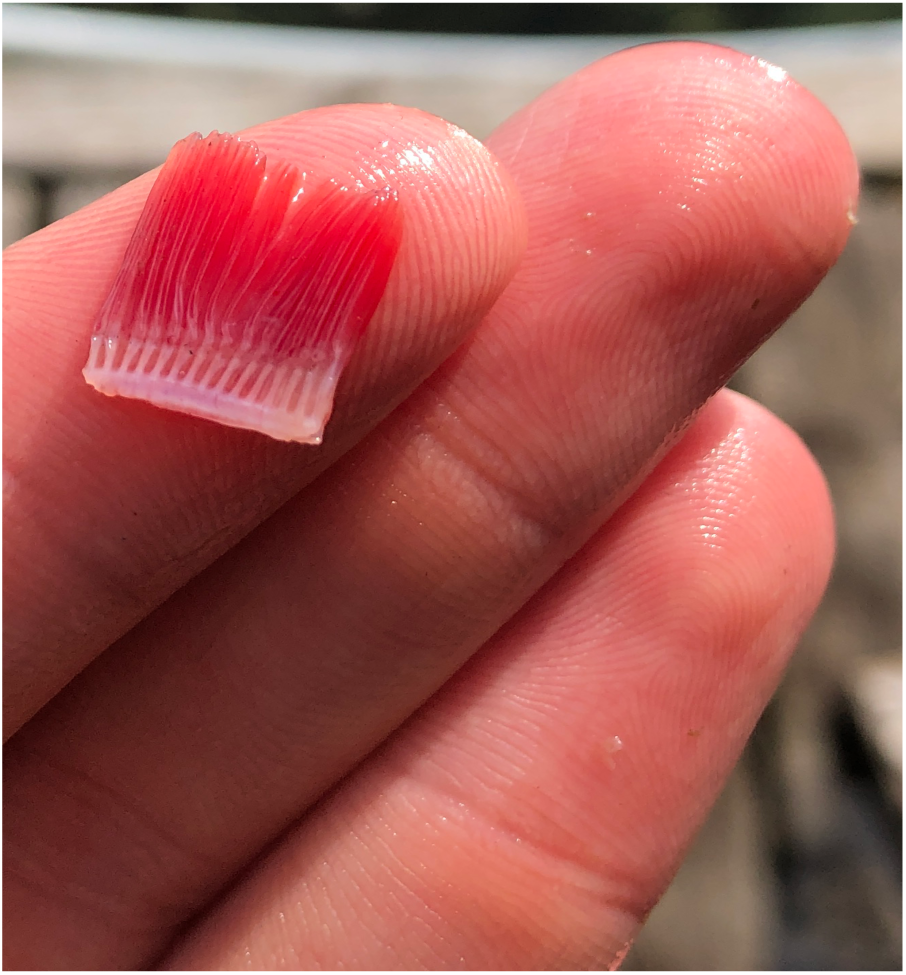
Image of the standardized 0.9 cm by 0.9 cm section of gill filaments removed from Northern Pike (*Esox lucius*) to simulate hooking injury.

Experiments were conducted at two different time periods. *Experiment 1* was conducted in May 2019 when water temperature ranged between 11-18 °C, whereas *Experiment 2* took place in August 2019 when water temperature ranged from 24-27 °C. *Experiment 1* had five treatment groups: 1) baseline, where the gill was not injured, and no carbonated beverages were used; 2) post injury, fish were held in lake water (pH = 6.71) without any other treatment; 3) post injury, carbonated lake water (pH = 4.37) poured over the gills; 4) post injury, Mountain Dew (pH = 3.27) poured over the gills; and 5) post injury, Coca-Cola (pH = 2.56) poured over the gills. During *Experiment 1*, beverage temperature ranged from 11-18 °C according to ambient air temperature. *Experiment 2* included baseline and control treatments (lake water pH = 6.71) as *Experiment 1*, and compared Mountain Dew (pH = 3.27) at ambient temperature (24-27 °C) with Mountain Dew (pH = 3.27) kept on ice (4-8 °C), to mimic an angler either not using or using a cooler, respectively, to hold beverages.

Following treatment fish were individually held in a cooler and visually monitored for 20 min to quantify: 1) time to bleeding cessation; 2) gill colour, which served as a proxy for blood loss; and 3) bleeding intensity. Bleeding cessation time was recorded as the time from gill injury until noticeable bleeding from the gill area stopped. Gill colour was assessed against a 20-point colour gradient, with bright red (20) at one end of the scale representing gills that were well-perfused with blood (most common), through progressively lighter shades of red to pink, to nearly white (1). Gill colour values were recorded 10 min and 20 min post-injury, as well as immediately before the gill injury occurred, serving as a reference. Relative bleeding intensity (BIN) was based on the following scale: 0, no bleeding; 1, minor bleeding, not obvious; 2, obviously bleeding, easily observed; and 3, intense bleeding, pulsatile blood flow. For all treatment groups other than the baseline group, we recorded bleeding intensity immediately before and after liquid was poured directly onto the wound while the fish was held in a water-filled trough. Additional bleeding intensity values were recorded at 3-min and 5-min post-injury. After 20 min in the cooler, the vigour and condition of the fish were recorded using reflex action mortality predictors (RAMP), and fish that were not moribund were released. Fish that were moribund were euthanized by cerebral percussion (3 fish in total; see below).

### Survival and Reflexes

The presence or absence of basic reflexes can be used to predict post-release mortality of fishes (Davis 2007; Raby et al. 2015). Our RAMP scoring system evaluated: 1) ability to maintain equilibrium; 2) reaction to grasping the tail (burst swimming); and 3) vestibular-ocular response (i.e. eye tracking) to determine whether a fish was impaired post-treatment and should be euthanized or released.

### Data Analysis

R Version 1.1.447, R Studio (R Core Team 2019) was used to conduct all statistical analyses. For both *Experiment 1* and *2*, bleeding time was compared among treatments using linear regression models and significant effects were assessed using Type 1 sum of squares. For bleeding intensity (BIN; ordinal scale from 0 to 3) and gill colour (ordinal scale from 1 to 20), ordinal logistic regression was applied to each time point at which BIN or gill colour was assessed. Full models included treatment, time, and their interaction as a fixed effect and individual as a random effect to account for repeated measures. Backward model selection was used to determine final model structure using Akaike Information Criterion (AIC). When best fit models included interactions, ordinal logistic regression models were fit within each sampling time period to assess differences among treatments. Statistical significance was accepted at α = 0.05 and, unless otherwise noted, all values are presented as means ± SEM.

## Results

For Experiment 1, 118 Northern Pike (50.9 ± 6.6 cm TL) were captured, while 38 Northern Pike (52.6 ± 6.7 cm TL, n = 38) were captured for *Experiment 2*. There were no significant differences in TL among the treatments for either experiment (*Experiment 1*, F_4_ = 0.641, p = 0.634; *Experiment 2*, F_3_ = 0.583, p = 0.629).

### Time to Bleeding Cessation

Time to bleeding cessation in *Experiment 1* ranged from 0 to 690 s (mean 193 ± 95 s) and was not significantly different among treatments (Fig. 2; F_3_ = 0.83, p = 0.48). For *Experiment 2*, time to bleeding cessation ranged from 0 to 193 s (mean 87 ± 40 s) and also not different among treatments (Fig. 2; F_2_ = 2.47, p = 0.10).

**Figure 2.**
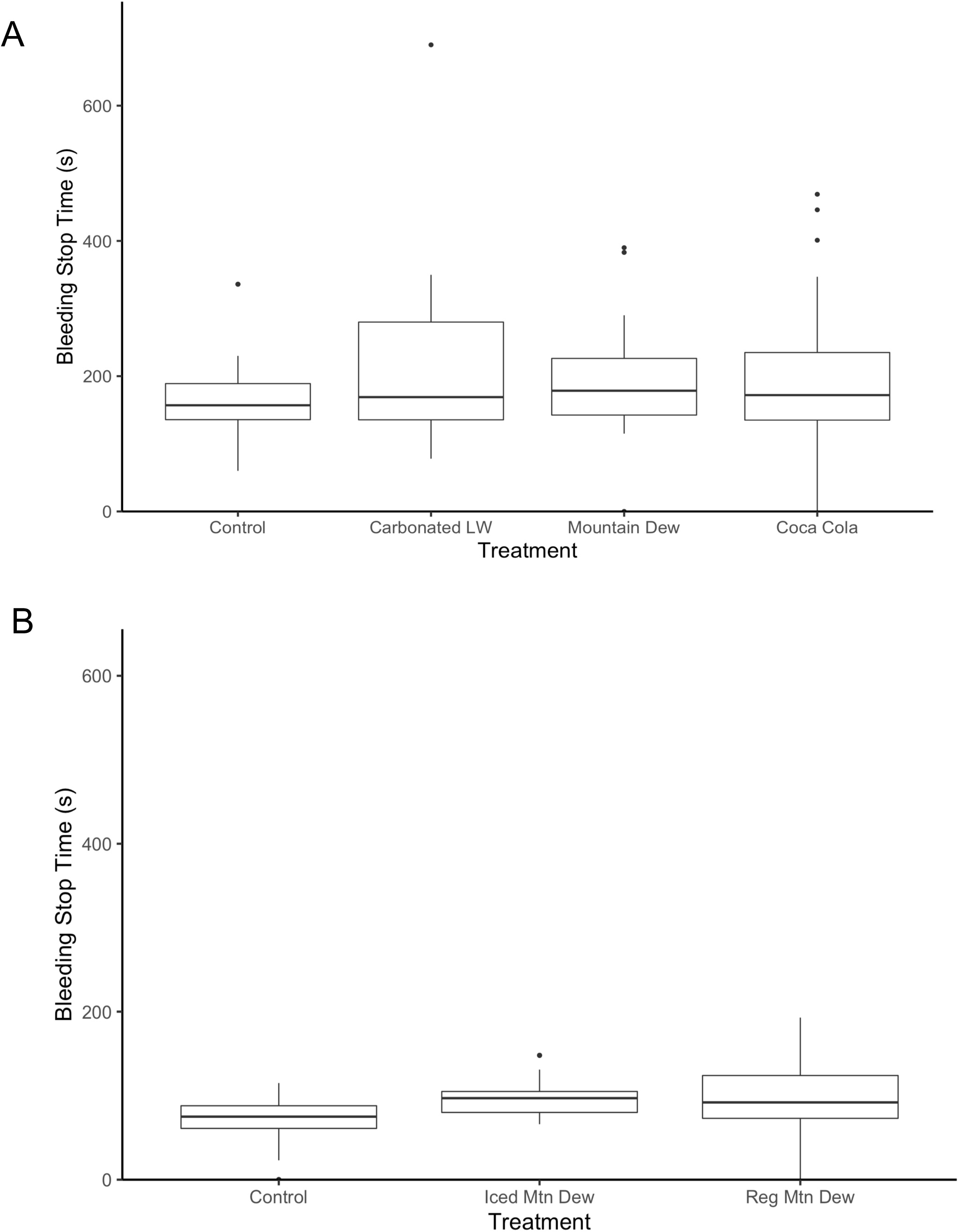
Time to cessation of bleeding following gill injury in Northern Pike in Experiment 1 (A) and Experiment 2 (B). In A, control is compared against carbonated lake water (Carbonated LW), Mountain Dew, and Coca Cola. In B) control is compared against Mountain Dew at 4-8°C (chilled Mountain Dew) and Mountain Dew at ambient (24-27°C (Reg Mountain Dew).

### Gill Colour Index

For *Experiment 1*, the best fitting model for gill colour index included a significant interaction between treatment and time (Table 1). Gill colour index did not differ among treatment groups prior to gill injury. Post injury, the baseline treatment (no injury) group exhibited significantly darker colour (higher score; Fig. 3) than all other treatment groups at both 10 min (t_126_ = 3.60, p < 0.001) and 20 min (t_127_= 5.03, p < 0.001). However, no significant differences were detected among the control group (immersion in lake water) and any group treated with a carbonated beverage (t < 5.03, p > 0.05; Table 2). In *Experiment 2* there was also a significant interaction between treatment and time (Table 1). The baseline treatment (no injury) had significantly higher colour score (darker colour) than all other groups both 10 min (t_31_ = 1.78, p <0.001) and 20 min (t_31_ = 3.62, p <0.001) post-injury (Fig. 4). There were no significant differences were detected among the control group and use of chilled Mountain Dew (t < −0.206, p >0.05). Fish had a significantly darker gill colour at 10 minutes when ambient temperature Mountain Dew was used in comparison to the control group (t_31_ = −2.309, p = 0.021; Table 2).

**Table 1:** Model selection outputs for ordinal logistic regression models. Full models included gill color as the response, the interaction between treatment and time as fixed effects, and individual as a random effect. Backward model selection was used to determine final model structure. Significant differences in model fit are highlighted by bold italic font.

| Experiment | Model Specification | AIC | Residual Deviation | Loglikelihood | P value | DF |
| --- | --- | --- | --- | --- | --- | --- |
| 1 | ***Treatment\*Time*** | ***2014.924*** | ***1954.924*** | ***25.575*** | ***<0.001*** | ***8*** |
|  | Treatment | 2086.274 | 2046.274 | -29.198 | 1.000 | 2 |
|  | ***Time*** | ***2053.076*** | ***2017.075*** | ***59.600*** | ***<0.001*** | ***2*** |
| 2 | ***Treatment\*Time*** | ***626.000*** | ***572.000*** | ***13.7312436*** | ***0.033*** | ***6*** |
|  | Treatment | 652.3781 | 614.378 | -0.0592199 | 1.000 | 1 |
|  | ***Time*** | ***650.3189*** | ***614.318*** | ***21.8639438*** | ***<0.001*** | ***2*** |

**Table 2:** Ordinal logistic regression model outputs for gill colour index with treatment as a predictor using control values as reference group for comparisons at individual time periods, 0, 10 and 20 minute. Significant differences are highlighted by bold italic font.

| Time | Comparisons | Value | Standard Error | T value | P-value |
| --- | --- | --- | --- | --- | --- |
| 0 Minutes | Carbonated Lake Water | 0.14346940 | 0.4523524 | 0.31716292 | 0.751 |
|  | Mountain Dew | 0.61190184 | 0.4527499 | 1.35152284 | 0.176 |
|  | Coca Cola | 0.14213540 | 0.4630298 | 0.30696813 | 0.758 |
|  | Baseline | 0.00598101 | 0.4948112 | 0.01208746 | 0.990 |
| 10 Minutes | Carbonated Lake Water | -0.40226948 | 0.4550232 | -0.88406369 | 0.376 |
|  | Mountain Dew | -0.19233292 | 0.4416917 | -0.43544612 | 0.663 |
|  | Coca Cola | -0.50898612 | 0.4646763 | -1.09535621 | 0.273 |
|  | **Baseline** | **1.78152855** | **0.4945991** | **3.60196492** | **<0.001** |
| 20 Minutes | Carbonated Lake Water | 0.1578421 | 0.4628604 | 0.3410145 | 0.733 |
|  | Mountain Dew | 0.6228802 | 0.4417568 | 1.4100071 | 0.158 |
|  | Coca Cola | -0.05273126 | 0.4547337 | -0.1159608 | 0.907 |
|  | **Baseline** | **2.500492** | **0.4969529** | **5.0316475** | **<0.001** |

Experiment 2 (n=38, df = 146)
| Time | Comparisons | Value | Standard Error | T value | P-value |
| --- | --- | --- | --- | --- | --- |
| 0 Minutes | Chilled Mountain Dew | -0.1457 | 0.7063 | -0.2063 | 0.8366 |
|  | Regular Mountain Dew | -0.4886 | 0.6783 | -0.7204 | 0.4713 |
|  | Baseline | 0.0071 | 0.7534 | 0.0094 | 0.9925 |
| 10 Minutes | Chilled Mountain Dew | -0.8734 | 0.7416 | -1.1777 | 0.2389 |
|  | **Regular Mountain Dew** | **-1.6813** | **0.7281** | **-2.3093** | **0.0209** |
|  | **Baseline** | **1.3084** | **0.7342** | **1.7821** | **0.0747** |
| 20 Minutes | Chilled Mountain Dew | -1.2230 | 0.7538 | -1.6225 | 0.1047 |
|  | Regular Mountain Dew | -0.2734 | 0.7305 | -0.3742 | 0.7082 |
|  | **Baseline** | **3.3882** | **0.9348** | **3.6243** | **<0.001** |

**Figure 3.**
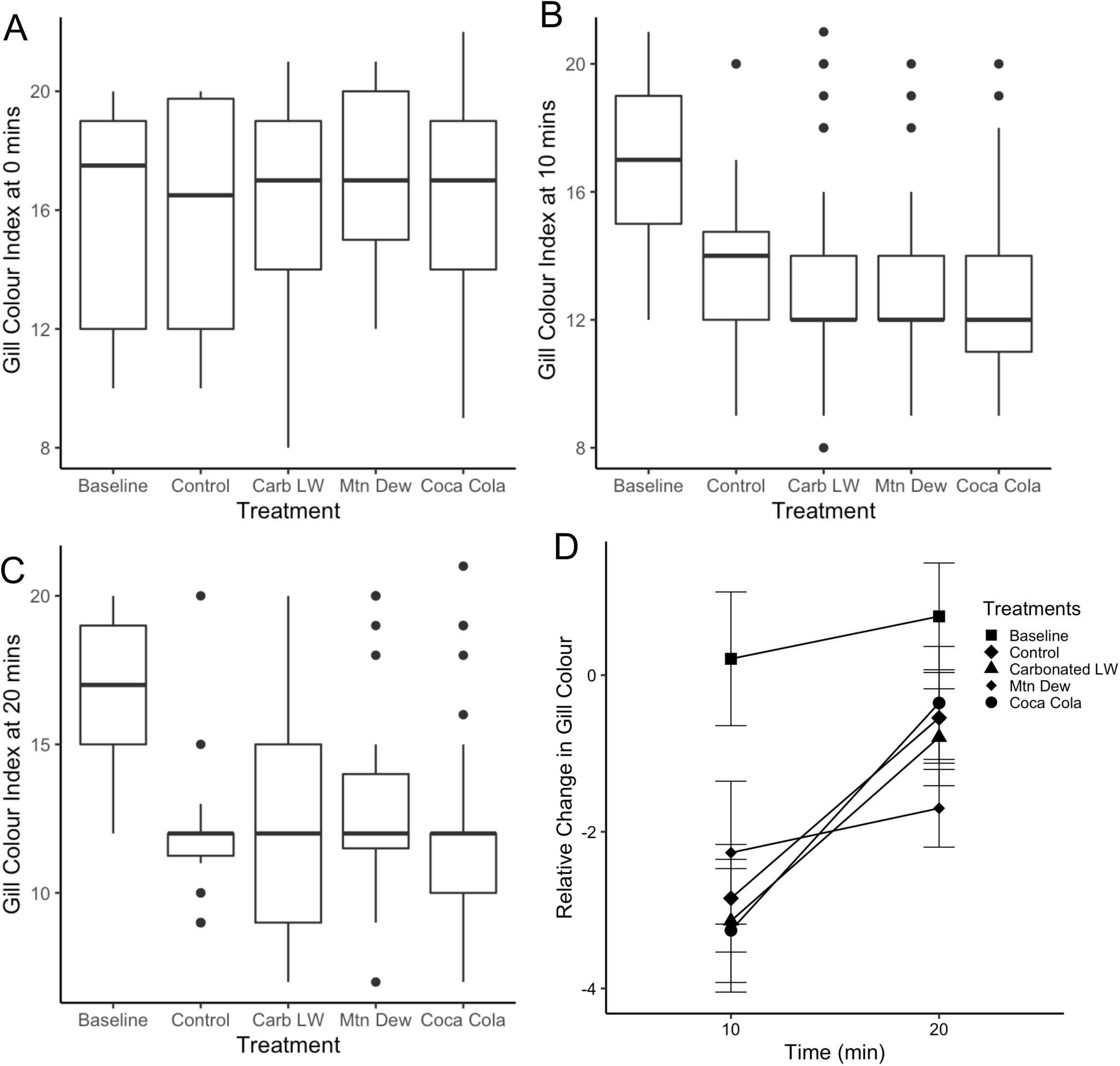
Gill Colour Index for *Experiment* 1 for 0 minutes (A), 10 minutes (B), 20 minutes (C) and the relative change from 0 to 10 minutes and 10 and 20 minutes (D). In all graphs control is compared against carbonated lake water (Carbonated LW), Mountain Dew, and Coca Cola.

**Figure 4.**
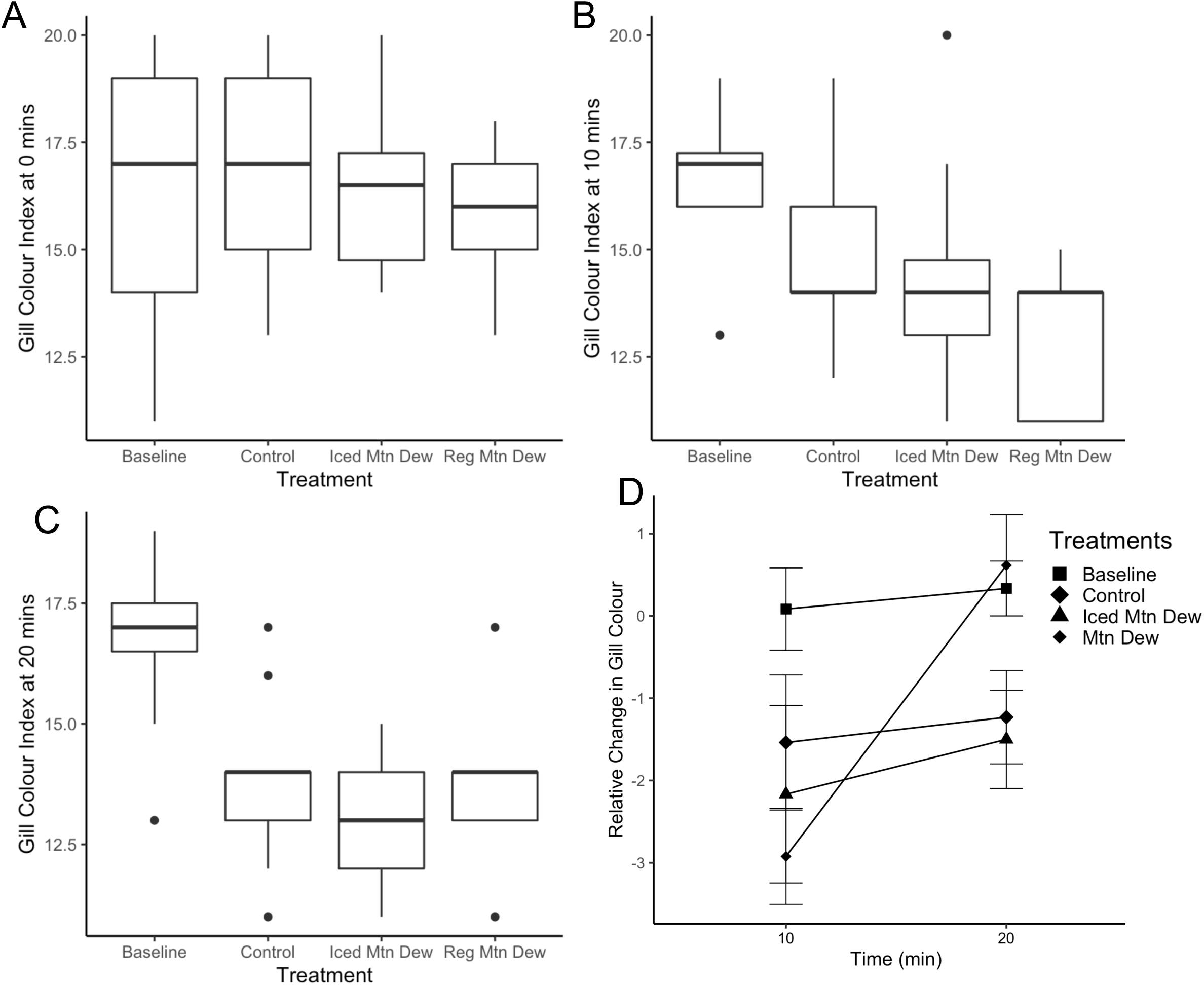
Gill Colour Index for Experiment 2 for 0 minutes (A), 10 minutes (B), 20 minutes (C) and the relative change from 0 to 10 minutes and 10 and 20 minutes (D). In all graphs control is compared against Mountain Dew at 4-8 °C (chilled Mountain Dew) and Mountain Dew at ambient temperature, 24-27 °C (Reg Mountain Dew).

### Bleeding Intensity Values

For *Experiment 1*, bleeding intensity was not different among treatments (BIN; Fig. 5A) as the best fitting model to the data included time as a fixed effect and individual as a random effect to account for repeated measures (Table 3). Similarly, treatment was not a significant contributing factor to variation in BIN in *Experiments 2* as the best fitting model to the data included time and individuals as independent predictors (Table 3).

**Table 3:** Model selection outputs for ordinal logistic regression models. Full models included bleeding intensity as the response, the interaction between treatment and time as fixed effects, and individual as a random effect. Backward model selection was used to determine final model structure. Significant differences in model fit are highlighted by bold italic font.

Experiment 1 (n = 118), Experiment 2 (n = 38)
| Experiment | Model Specification | AIC | Residual Deviation | Loglikelihood | P value | DF |
| --- | --- | --- | --- | --- | --- | --- |
| 1 | Treatment*Time | 531.4823 | 495.4823 | 9.473 | 0.394 | 9 |
|  | Treatment | 1030.8998 | 1018.8998 | 0.059 | 0.996 | 3 |
|  | **Time** | **520.6760** | **508.6760** | **510.223** | **<0.001** | **1** |
| 2 | Treatment*Time | 230.3625 | 202.3625 | 1.7381 | 0.942 | 6 |
|  | Treatment | 361.4302 | 351.4302 | 0.600 | 0.741 | 2 |
|  | **Time** | **217.4912** | **205.4912** | **145.939** | **<0.001** | **1** |

**Figure 5.**
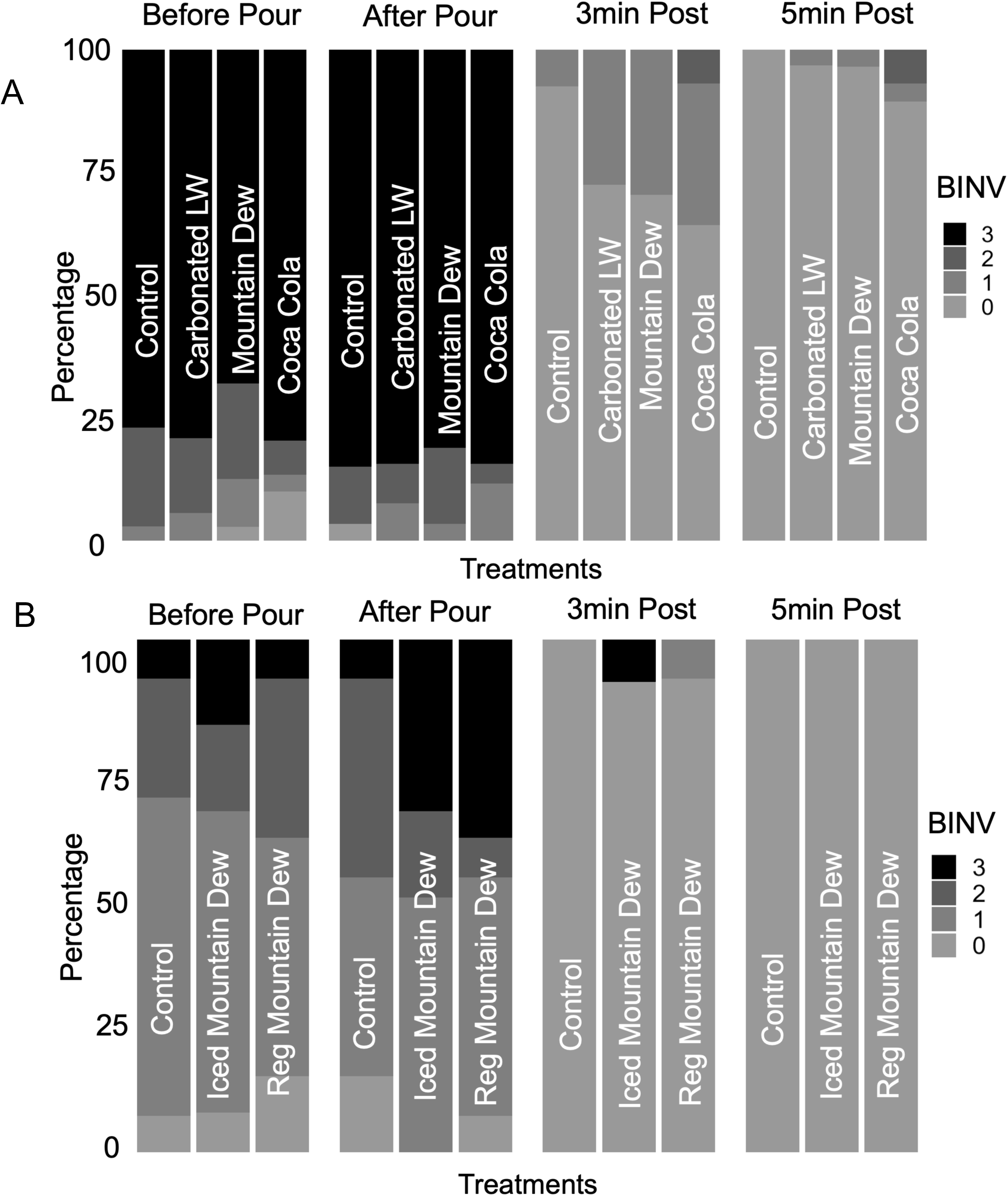
Bleeding intensity values following gill injury in Northern Pike in Experiment 1 (A) and Experiment 2 (B). Figures show the bleeding intensity values before the pour of any substance, after the cut (Before Pour), after the pour of a substance (After), after 3 minutes (3min Post) and 5 minutes after the substance was poured over the gills (5min Post). In A, control is compared against carbonated lake water (Carbonated LW), Mountain Dew, and Coca Cola. In B) control is compared against Mountain Dew at 4-8°C (chilled Mountain Dew) and Mountain Dew at ambient (24-27°C (Reg Mountain Dew).

### Survival and Reflexes

In *Experiment 1*, no significant effect of treatment group on RAMP score was detected (F_4_ = 1.008, p = 0.405). In *Experiment 2*, a significant treatment effect was detected (F_3_ = 4.002, p = 0.013), with fish subjected to chilled Mountain Dew exhibiting significantly higher impairment than the baseline group (t_46_ = −3.43, p = 0.001). Only 3 fish were euthanized owing to low RAMP score (2 fish in the carbonated lake water group and 1 in the Coca Cola group during *Experiment 1*).

## Discussion

The main claim of those in the recreational angling community promoting the use of carbonated beverages is that this practice decreases the duration bleeding related to hooking injury, particularly of the gills (Pyzer 2019). However, our study found no differences in the time to cessation of bleeding, gill colour (which was used as an index of blood loss), or bleeding intensity among three carbonated beverages poured over bleeding gills, or between the carbonated beverages and a control group. Differences were only detected between the baseline group where no simulated hooking injury occurred, and all other groups in which injury and bleeding happened. These findings provide direct scientific evidence that the use of carbonated beverages does not curtail bleeding from gills, which is counter to anecdotal observations made by recreational anglers that use this technique. Overall, our study debunks the assertion that pouring carbonated soft drinks on the gills of injured via recreational angling is a best practice for C&R at least in the context studied here.

Although some online forums assert that exposure to CO_2_ from carbonated beverages would be beneficial for reducing or even eliminating bleeding from the fills of fish, scientific studies suggest that exposure to high environmental CO_2_ has negative effects on fishes (Perry and Abdallah 2012; Kaya et al. 2016; Tresguerres et al. 2019). For example, CO_2_ excretion and acid-base regulation are impacted by acute exposure to elevated CO_2_ and high water CO_2_ triggers cardiorespiratory reflexes such as hyperventilation and bradycardia (Taylor 1988; Gilmour 2001; Perry and McKendry 2001; Claiborne et al. 2002; Perry and Reid 2002; Gilmour et al. 2005; Gilmour 2012). As such, the physiological effects of acute CO_2_ exposure imply that carbonated beverages are unlikely to benefit the well-being of fishes.

Hypoxia-induced bradycardia in combination with CO_2_-induced bradycardia may present a perceived positive effect when carbonated beverages are initially poured on fish gills that are bleeding. Many anglers lift fish out of the water once landed, exposing them to air, promoting hypoxia (Cooke and Sneddon 2007), which can induce bradycardia (Randall 1982; Farrell 2007). This effect of air exposure on fish has been well characterized in angled fish (Cooke et al. 2002; 2003; Furimsky et al., 2003). Therefore, when fish are removed from the water so that carbonated beverages can be used on the gills, the perception that bleeding has ceased is possible, but the effect is largely driven by hypoxia-induced decreases in cardiovascular output (Reid and Perry 2003; Perry and Desforges 2006). For our study, fish were held horizontally in a water-filled trough such that the gills were constantly submerged in well-oxygenated water. This approach should have limited hypoxia-induced bradycardia allowing effects of the carbonated beverages to be detected. Because we used white coolers, CO_2_-induced bradycardia effects were apparent. Fish would not bleed for >30 s, followed by blood spurting from the gills. If a fish were to be released by an angler or held in the water, this bleeding might not be apparent, and could account for reports of carbonated beverages stopping blood loss (for a short period).

Independent of the effects of carbonated beverages on bleeding, these acidic solutions may have damaging effects on fish gill tissues. Low pH water has been shown to cause a general inflammatory response, an increase in mucous production, and alterations in the structural morphology of the gills in teleost fish (Meyer et al. 2009). Although many of these studies use more chronic exposures (i.e. > 24 h) than used in our study, acute exposure may still have negative effects on fish (Meyer et al. 2009), particularly on ion and acid-base regulation (Wright and Wood 2009). Interestingly, Northern Pike are relatively tolerant of environmental acidification and have reported to survive in water of pH 3.5 (Zaprudnova et al. 2015), which may allow them to cope with an acute acid exposure.

Further work should address how acute instances of branchial acid exposure can affect ion and acid-base status in this context to fully appreciate the biological consequences of using carbonated beverages in an angling setting. Also, given that we limited our observations for 20 min during our study, we did not assess more chronic physiological can behavioral effects, as well as post-release mortality. Although Northern Pike are regarded as being relatively robust to hooking injury (Arlinghaus et al. 2008), and that we observed little immediate or short-term mortality in fish that were actually injured during capture for this study, the longer-term consequences of pouring chemicals associated with carbonated sodas remains unknown. Future work involving telemetry or net pens would be useful for understanding longer-term consequences of this carbonated beverage technique. Lastly, our study used a simulated and standardized size of gill injury to a single species of fish, so further investigation is are needed to determine whether the magnitude of injury or species-specific differences would produce different results.

Overall, we provide the first evidence that counters the growing popularity of using carbonated beverages to stop bleeding in angling-caught fish. Our observations made during the 20 min holding period versus immediately release as many anglers do after they pour carbonated beverages on injured gills, sheds light on the potential perception of the curtailment of bleeding anglers witness, especially if the combination of air exposure and CO_2_ cause an immediate and severe bradycardia. We found no benefit or disbenefit with pouring carbonated beverages over the gills of Northern Pike, but it is possible that there are longer term impacts. Similarly, our findings are specific to the context studied here. As such, it is possible that carbonated beverages could provide benefit or harm when used in other contexts, such as with other species (e.g., salmonids, muskellunge), using other beverages, or applying the beverages in other ways (e.g., holding them in air for longer after applying beverage). This study contributes to the growing body of literature that emphasizes the need for anglers and fisheries scientists to work collaboratively to ensure that best practices being employed truly benefit fish (Brownscombe et al. 2017).

## Acknowledgements

We acknowledge John Anderson from the Ottawa Musky Factory for encouraging us to pursue this research. We extend our gratitude to Queen’s University Biological Station for serving as a base for this research. Adam Williamson, Jon Kulbeka, Brooke Etherington and the many other researchers than helped make this project possible. Ben Hlina provided input on statistical analysis and figures. The document was kindly reviewed by Katherine Gilmour. Funding was provided by Muskies Canada and The Anderson Family Foundation with additional support from the Natural Sciences and Engineering Research Council of Canada and the Canada Research Chairs Program. All research was conducted in accordance with the guidelines of the Canadian Council on Animal Care as administered by Carleton University (Protocol 110558).

